# Effects of Pesticide Exposure on Neuroinflammation and Microglial Gene Expression: Relevance to Mechanisms of Alzheimer’s Disease Risk

**DOI:** 10.1101/2025.02.14.638293

**Authors:** Isha Mhatre-Winters, Aseel Eid, Nicole Blum, Yoonhee Han, Ferass M. Sammoura, Long-Jun Wu, Jason R. Richardson

## Abstract

**Background:** Alzheimer’s disease (AD) is characterized by the presence of amyloid-β plaques, neurofibrillary tangles, and neuroinflammation. Previously, we reported serum levels of dichlorodiphenyldichloroethylene (DDE), the primary metabolite of the pesticide dichlorodiphenyltrichloroethane (DDT), were significantly higher in AD patients compared to age-matched controls and that DDT exposure worsened AD pathology in animal models.

**Objective:** Here, we investigated the effect of DDT on neuroinflammation in primary mouse microglia (PMG) and C57BL/6J mice.

**Methods:** Effects of DDT on inflammation and disease-associated microglia were determined in primary mouse microglia and C57BL/6J mice.

**Results:** PMG exposed to DDT (0.5-5.0 µM) elicited a ∼2-3-fold increase in *Il-1b* mRNA levels, with similar concentration-dependent upregulation in *Il-6, Nos2,* and *Tnfa*. These effects were blocked by the sodium channel antagonist tetrodotoxin, demonstrating the role of DDT-microglial sodium channel interactions in mediating this response. Additionally, NOS2 protein levels increased by ∼1.5-2-fold, while TNFa was elevated by 2-4-fold. C57BL/6J male and female mice exposed to DDT (30 mg/kg) demonstrated significantly increased mRNA levels of *Nos2*, *Il-1b*, and *Il-6* in the frontal cortex (1.5-2.3-fold), and *Nos2*, *Il-1b,* and *Tnfa* (1.5-1.8-fold) in the hippocampus. Furthermore, microglial homeostatic genes, *Cx3cr1*, *P2ry12,* and *Tmem119*, were downregulated, while stage 1 disease-associated microglia genes were upregulated both *in vitro* and *in vivo*. Notably, *Apoe* and *Trem2* were only upregulated in the frontal cortex and hippocampus of females.

**Conclusion:** These data indicate that DDT increases neuroinflammation, which may result from direct actions of DDT on microglia, providing a novel pathway by which DDT may contribute to AD risk.

## Introduction

Neuroinflammation is implicated in the progression and pathogenesis of several neurodegenerative diseases, including Alzheimer’s Disease (AD). While AD research has traditionally focused on mechanisms involved in extracellular amyloid plaques and intracellular neurofibrillary tangle formation, recent evidence from genome-wide association studies (GWAS) has revealed several risk loci in genes related to the immune system [1-6]. These findings have further strengthened the importance of activated glia in the pathology and progression of the multifaceted disease [7, 8].

Neuroinflammation is a concerted process predominantly initiated by glial cells in the brain, specifically microglia and astrocytes. Glial cells, when stimulated due to disease, injury, infection, or stress, can produce inflammatory mediators such as cytokines, chemokines, reactive oxygen species, and secondary messengers [9, 10]. Microglial activation involves increased synthesis and release of cytokines and chemokines, which are regulated by cell surface receptors and ion channels. Depending on the response of microglia, early studies classified microglia into 2 states: M1 – proinflammatory or neurotoxic and M2 state – anti-inflammatory [11]. However, these binary categories did not explain specific gene expression profiles of reactive microglia, specifically those observed in neurodegenerative diseases such as AD [12-14]. Recent single-cell and single-nucleus sequencing studies identified and characterized a dynamic state of microglia called disease-associated microglia (DAM) [15, 16]. This microglial phenotype was shown to cluster around amyloid-β plaques in AD mouse models and in human AD postmortem brains as well as downregulate homeostatic microglial genes and upregulate genes involved in lysosomal, phagocytic, and lipid metabolism pathways [1, 17, 18].

Microglia have been shown to express voltage-gated sodium channels (VGSC), which regulate multiple functions, including migration, proliferation, morphological transformation, phagocytosis, and secretion of cytokines [19-22]. Previous studies have shown a dose-dependent increase in the accumulation of sodium ions (Na^+^) and an inward current through VGSC after exposure to the endotoxin lipopolysaccharide (LPS) [23]. This increase was inhibited by the VGSC blocker tetrodotoxin (TTX), suggesting that LPS activates microglia via an interaction with the VGSC [23]. Consistently, in multiple models following LPS exposure, TTX inhibited secretion and expression of pro-inflammatory cytokines, including TNFa, IL-1b, NOS2, and production of reactive oxygen species [24].

VGSCs are the primary target of both dichlorodiphenyltrichloroethane (DDT) and pyrethroid insecticides. Both classes of pesticides target VGSCs by delaying their closure, leading to hyperactivity of the nervouse system and, at high enough concentrations, death [25-28]. Previously, we reported that the pyrethroid insecticides, deltamethrin and permethrin, increased the secretion of the pro-inflammatory cytokine TNFa from microglia [29]. This was accompanied by a rapid influx of Na^+^ into the cell, which can be partially inhibited by the inclusion of TTX [29]. Here, we investigated the effects of DDT on neuroinflammation, a primary contributor to many neurodegenerative diseases, including AD. The data reveal direct actions of DDT on primary mouse microglia (PMG), but no effects on primary mouse astrocytes (PMA), suggesting the effects of DDT on cytokine expression and release are specific to microglial cells. Direct DDT exposure increased pro-inflammatory mRNA and proteins in the PMG, and this increase was blocked by the inclusion of TTX, demonstrating the requirement of interaction of DDT with VGSCs. C57BL/6J mice exposed to a single dose of 30 mg/kg DDT demonstrated significantly increased expression of pro-inflammatory cytokines in the frontal cortex and hippocampus. Lastly, we show that exposure to DDT, *in vitro* and *in vivo*, results in a transition of homeostatic microglia to disease-associated microglial (DAM) phenotype, a prominent feature observed in AD brains. Collectively, these data indicate that exposure to DDT increases neuroinflammation in C57BL6/J mice and is involved in directly activating microglia, providing a novel pathway by which DDT exposure may contribute to AD risk.

## Methods

### Animals

Animal studies were reviewed and approved by the Institutional Review Board (or Ethics Committee) of Florida International University (approval code: IACUC-21-047). All studies complied with the Animal Research: Reporting of In Vivo Experiments (ARRIVE) guidelines [30] and were carried out in accordance with the National Institutes of Health (NIH) *Guide for the Care and Use of Laboratory Animals* [31] and approved by the Institutional Animal Care and Use Committee of Florida International University. Male and female C57BL/6J mice (Jackson Laboratory, Bar Harbor, ME) were bred in-house and housed in a 12-hour light-dark cycle at 22±2°C and 50±10% of relative humidity. Mice had access to food (standard chow) and water *ad libitum*. Breeders were paired at 8-10 weeks of age to obtain mouse pups for primary cultures. Each litter was used as individual isolation for experiments.

### Reagents

All cell culture reagents were obtained from Invitrogen. Chemical reagents were purchased from Sigma Aldrich, except for *p,p’-*DDT (Chem Service, Cat. No. CN-10876, 99.20 % pure) and TTX (Alomone Labs, Cat. No. T-500).

### Primary mouse microglia (PMG) and primary mouse astrocyte (PMA) isolation

PMG and PMA were isolated using postnatal 0-2 day old C57BL/6J pups (mixed-sex) and maintained using well-established methods [32, 33]. Briefly, whole brains were collected, trypsinized, and triturated in growth media and passed through a 70 μm strainer. Cultures consisting of mixed glial cell types were grown in flasks for 15 days, followed by separating microglia using a CD11b positive selection kit (STEMCELL Technologies, Cat. No. 18970). The negative fraction containing PMA was grown in flasks for 4-5 days before seeding for experiments. Following isolation, cell purity for both PMG and PMA was assessed by immunocytochemistry (described below), and purity of greater than >97% were consistently obtained for both **(Figure 1 A - D)**. PMG were grown in Dulbecco’s Modified Eagle’s Medium/F12 (DMEM/F12) supplemented with 10% heat-inactivated fetal bovine serum (HI-FBS), 2mM L-glutamine, 1mM sodium pyruvate, 100 µM non-essential amino acids, 1% Penicillin/Streptomycin and 2.5 μg/mL Plasmocin™. PMA were grown in DMEM supplemented with 10% HI-FBS, 2mM L-glutamine, 1mM sodium pyruvate, 1% Penicillin/Streptomycin, and 2.5 μg/mL Plasmocin™.

**Figure 1:**
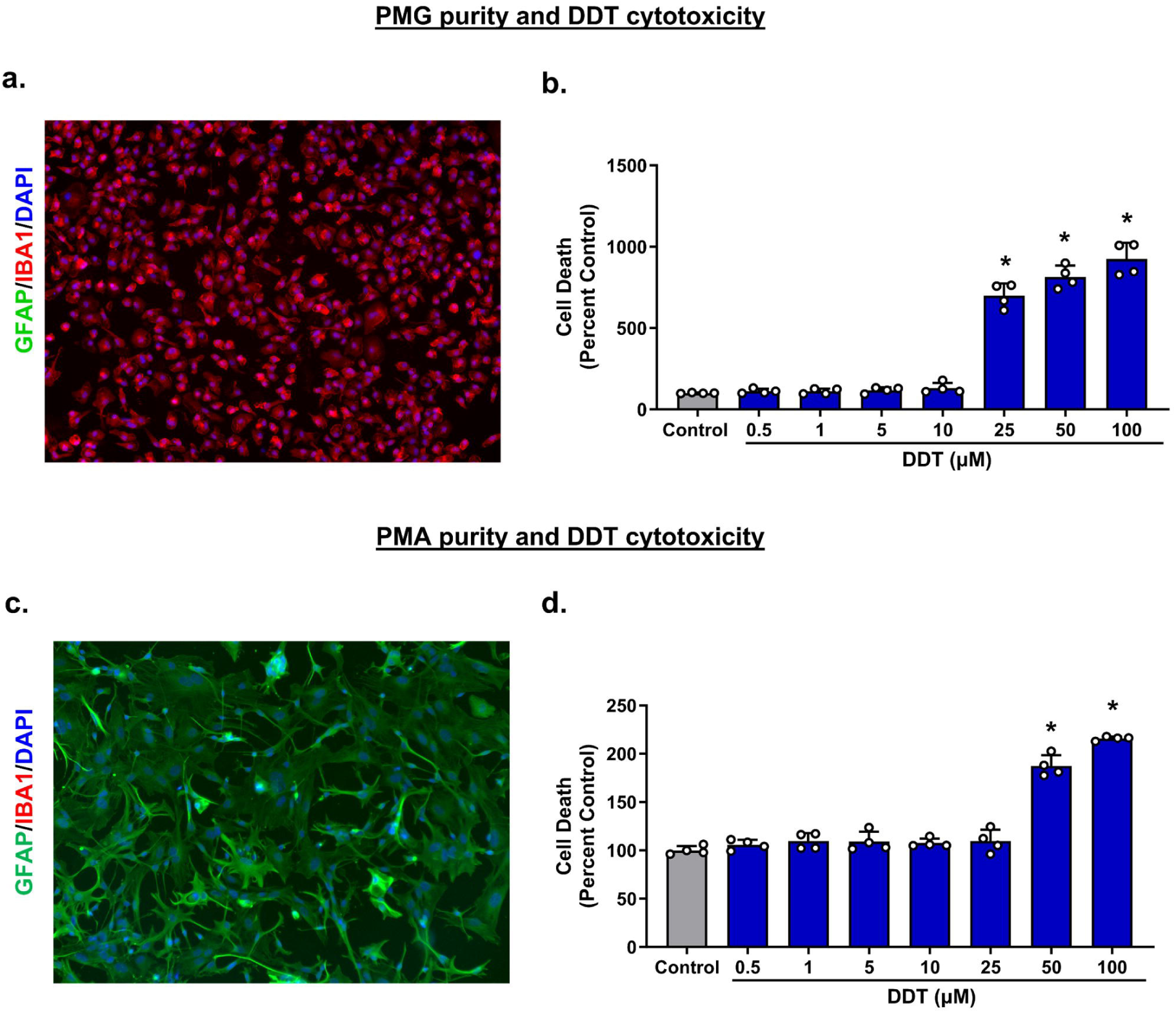
Cytotoxicity of purified PMG and PMA cultures following DDT exposure. Representative staining of isolated PMG (20x) (a) and SYTOX® Green assay for increased concentrations of DDT exposure on PMG (b). Representative staining of isolated PMA (c) and SYTOX® Green assay on PMA (d). n=4 independent isolations consisting of 3 technical replicates each. Data are presented as means and error bars as ± SD. Statistical significance was determined using a One-way ANOVA. * indicates p<0.05; scale bar= 31 pM.

### In vitro exposure

PMG and PMA were seeded into plates and treated after 24 hours. The *p,p’-*DDT stock was prepared in dimethyl sulfoxide (DMSO) and TTX in water. Dilutions from the stock were made in PMG or PMA growth media supplemented with 2% HI-FBS. Control groups were treated with 0.1% DMSO in the treatment medium. Cells were treated for 6 and 24 hours to assess gene expression and protein measurements, respectively. For treatments including TTX, cells were pre-treated for 15 min with 1 μM TTX for PMG and 10 μM TTX for PMA, followed by the addition of respective treatments.

### Cytotoxicity assay

PMG and PMA were plated in black-walled clear bottom 96-well plates at 80,000 and 30,000 cells/well, respectively. Following treatment with *p*,*p*’-DDT (0.5-100 µM) or 0.1% DMSO (vehicle control) for 24 hours, cell death was evaluated with SYTOX Green Nucleic Acid Stain following the manufacturer’s instructions (Invitrogen, Cat. No. S7020).

### Immunocytochemistry (ICC) and ICC quantification

PMG and PMA were seeded into 96-well plates at 80,000 and 30,000 cells/well, respectively. Following treatment, cells were fixed on slides with 10% formalin for 15 minutes, washed with PBS, and ICC was performed as previously described [32]. Briefly, cells were permeabilized with PBS + 0.5% Triton X-100 and blocked for 1 hour in 2% BSA solution at room temperature. Cells were then incubated overnight in 2% BSA with the following primary antibodies, Gfap (Glial fibrillary acidic protein; 1:1000, Abcam, Cat. No. ab4674) and Iba1 (Ionized calcium-binding adapter molecule; 1:1000, Wako chemicals, Cat. No. 019-19741) at 4°C to assess for purity. The following day, cells were washed with PBS and incubated with secondary antibodies from appropriate species (Alexa Fluor® 1:1500) for 1 hour at room temperature. Cells were assessed for purity in culture by imaging 3 to 4 random fields per well in a 96-well plate using the 10× objective on the Keyence BZ-X810 microscope. Cells positive for Iba1 (microglia) or Gfap (astrocytes) were counted against the total number of cells (positive for 4′,6-diamidino-2-phenylindole (DAPI)) per field using the BZ-X800 Analyzer software, and the average was determined as percent purity. Cultures consisting of >98% purity were consistently obtained.

PMG were also incubated overnight with Iba1 and Nos2 (1:500, Santa Cruz Biotechnology, Cat. No. sc-7271) or Iba1 with Tnfa (1:100, Abcam, Cat. No. ab1793) at 4°C. The next day, cells were washed with PBS and incubated with secondary antibodies from appropriate species (Alexa Fluor® 1:1500) for 1 hour at room temperature. The specificity of staining was confirmed by omitting primary or secondary antibodies. Protein levels of Tnfα and Nos2 in PMG were quantified using fluorescence intensity by integrated density analysis (Image J Software version 1.51j8). 5-6 fields per treatment were imaged using a Leica DMi8 SPE II confocal microscope using a 63x oil lens. The average intensity per cell was calculated for each treatment and compared to control, and data are represented as fluorescence intensity relative to control.

### In vivo exposure

Male and female C57BL/6J mice were randomly assigned to either DDT exposure or control groups. Mice between 17-19 weeks of age received either a single dose of 30 mg/kg of DDT dissolved in corn oil or corn oil (vehicle control) by oral gavage (n=4-6 per group). This dose is 10 times lower than the serious lowest observed adverse effect level (sLOAEL; 200-300 mg/kg/day) [34] reported in mice after receiving a single dose of *p*,*p*’-DDT [35]. Thus, this dosage does not cause overt toxicity to animals and was used to assess an acute inflammatory response to *p*,*p*’-DDT. Animals were monitored throughout the study and sacrificed 24 hours post-exposure by isoflurane administration followed by decapitation. The hippocampus and frontal cortex were dissected from the right hemisphere, frozen in liquid nitrogen, and stored at - 80°C until further processing.

### RT-qPCR

PMG and PMA were seeded into 24-well plates at 400,000 and 70,000 cells/well, respectively. RNA was isolated from PMG, PMA, and tissue (hippocampus and frontal cortex), using Trizol Reagent (Thermo Fisher Scientific, Cat. No. 15-596-018), following the manufacturer’s instructions. cDNA was synthesized using the All-in-One cDNA Synthesis Supermix (Bimake, Cat. No. B24408). Primer sets were verified with NCBI BLAST and ordered from Thermo Fisher Scientific. PCR reactions were performed using SYBR green (Bimake, Cat. No. B21203), and samples were run in duplicate on the QuantStudio 6 Flex System (Applied Biosystems). A list of genes and primer sequences used in qPCR reactions are presented in Supplemental Table 1. Gene expression was normalized to *Rpl13a* using the 2^-ΔΔCT^ method [36].

### Statistics and Software

Statistical analysis was performed using Prism 7.04 (GraphPad Software, San Diego, CA). Data were assessed for normality by the Shapiro–Wilk normality test. For the *in vitro* experiments, one-way or two-way analysis of variance (ANOVA) was used where appropriate and indicated in the figure legend and results section. For analyzing the *in vivo* data, sexes were analyzed seprately *a priori*, as estrogen and estradiol alter VGSC expression [37, 38]. Student’s t-test was used to compare the DDT-exposed group to the control group in the *in vivo* studies. Statistical significance was considered at p<0.05. All data are presented as means and error bars as ± standard deviation.

## Results

### DDT exposure increases mRNA and protein levels of pro-inflammatory cytokines

In these studies, we isolated and utilized purified cultures of PMG **(Fig 1 a)** and PMA **(Fig 1 c)**. In all *in vitro* experiments, the concentrations of DDT treatments were below the level that induced overt toxicity to the indicated cell type **(Fig 1b and d)**. Recent evidence suggests that both microglia and astrocytes are functionally homogenous and exist as differential phenotypes throughout the brain [39, 40]. These phenotypes range from resting state to pro-inflammatory phenotype, expressing cytokines including *Il-1b*, *Il-6*, *Tnfa*, *Ifng*, and *Nos2,* to the anti-inflammatory and regenerative phenotype [41, 42]. We first sought to assess whether DDT induced a pro-inflammatory phenotype in microglia. To this end, PMG were treated with 0.5, 1, and 5 μM of DDT, and mRNA levels of pro-inflammatory cytokines were analyzed using one-way ANOVA. A significant effect of treatment for *Il-1b* (F_3,12_ =23.66, p<0.0001), *Il-6* (F_3,12_ =12.63, p=0.0005), *Nos2* (F_3,12_ =19.78, p<0.0001), and *Tnfa* (F_3,12_ =26.19, p<0.0001) was observed **(Figure 2 a-d)**. Following exposure to 0.5 μM DDT, one-way ANOVA and Dunnett’s multiple comparisons test indicated an approximate 2-fold increase in gene expression of pro-inflammatory factors such as *Il-1b* (p=0.007), *Nos2* (p=0.006) and *Tnfa* (p=0.0006). After exposure to 1 μM DDT, we measured a 2.8-fold increase in *Il-1b* (p=0.0002), a 1.6-fold increase in *Il-6* (p=0.002), a 2.6-fold increase in *Nos2* (p=0.0001) and *Tnfa* (p<0.0001). Lastly, after exposure to 5 μM DDT, *Il-1b* was significantly increased by 3.4-fold (p<0.0001), *Il-6* increased by 1.7-fold (p=0.004), while both *Nos2* and *Tnfa* increased by 2.9-fold (p<0.0001). In contrast, in the PMA, at all treated concentrations of DDT, no significant increase in pro-inflammatory cytokine expression was observed **(Figure S1)**.

**Figure 2:**
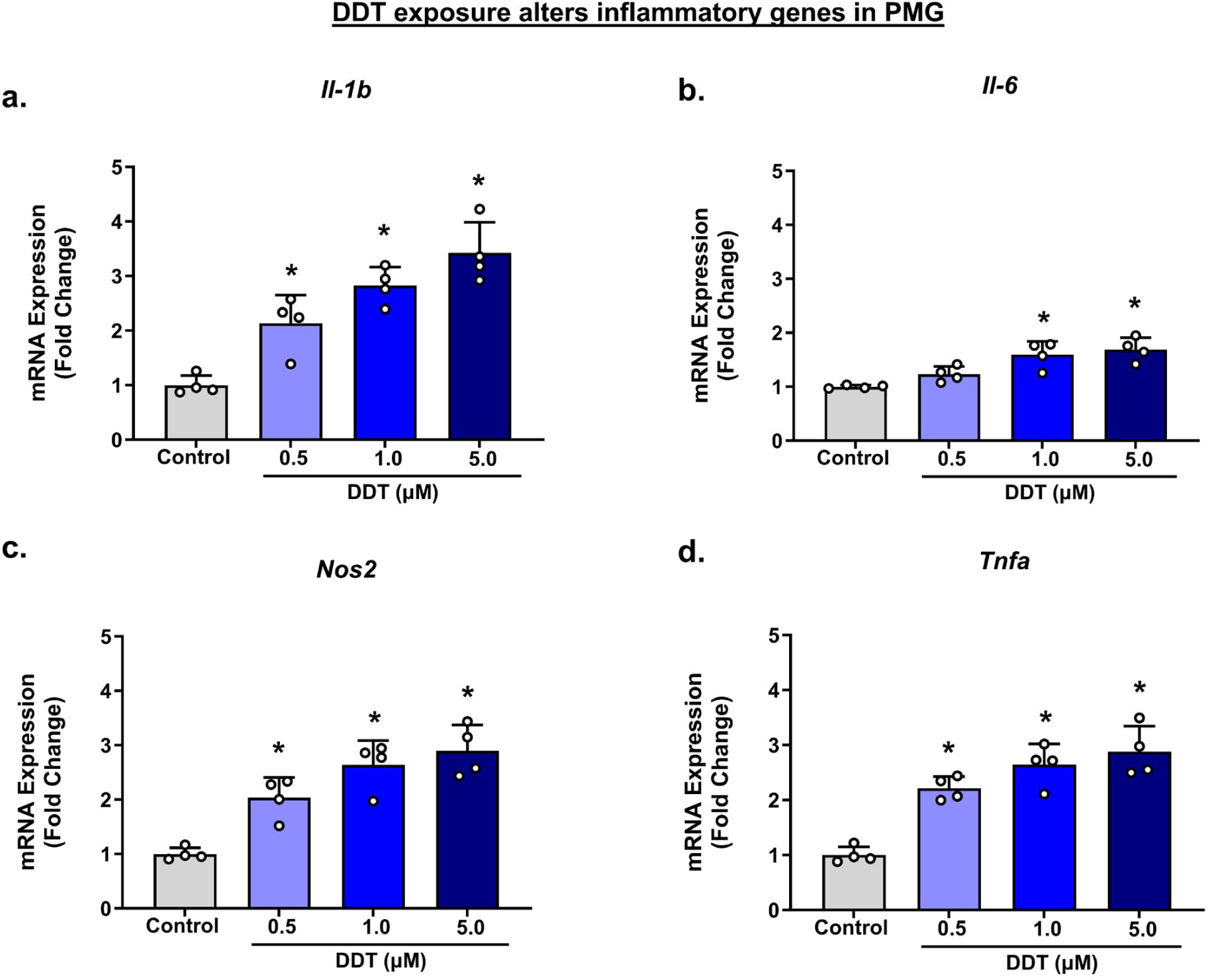
DDT exposure increases gene expression of inflammatory factors in PMG. mRNA levels of inflammatory genes in PMG exposed to DDT for 6 h (a - d). n=4 independent isolations consisting of 3 technical replicates each. Data are presented as means and error bars as ± SD. Statistical significance was determined using a One-way ANOVA. * indicates p<0.05.

To examine whether this increase in gene expression in PMG was accompanied by protein changes in Tnfa and Nos2 levels in the cell, we performed ICC following 24-hour DDT treatment to PMG. One-way ANOVA revealed a significant effect of DDT exposure on PMG for Nos2 protein levels by ICC (F_3,12_ =24.71, p<0.0001), with an increase of 39% at 0.5 μM (p=0.008), 69% at 1.0 μM (p<0.0001), and 84% at 5.0 μM (p<0.0001) relative to control **(Figure 3 a and b)**. For TNFa protein levels, similar results of DDT exposure were measured by one-way ANOVA (F_3,12_ =36.75, p<0.0001) with a significant increase of 48% at 0.5 µM (p=0.024), 185% at 1.0 µM (p=0.0001) and 240% at 5.0 μM (p<0.0001) relative to control **(Figure 3 c and d)**. These results collectively demonstrate that DDT directly activates microglia, and not astrocytes, and is associated with the upregulation of pro-inflammatory gene expression and proteins *in vitro*.

**Figure 3:**
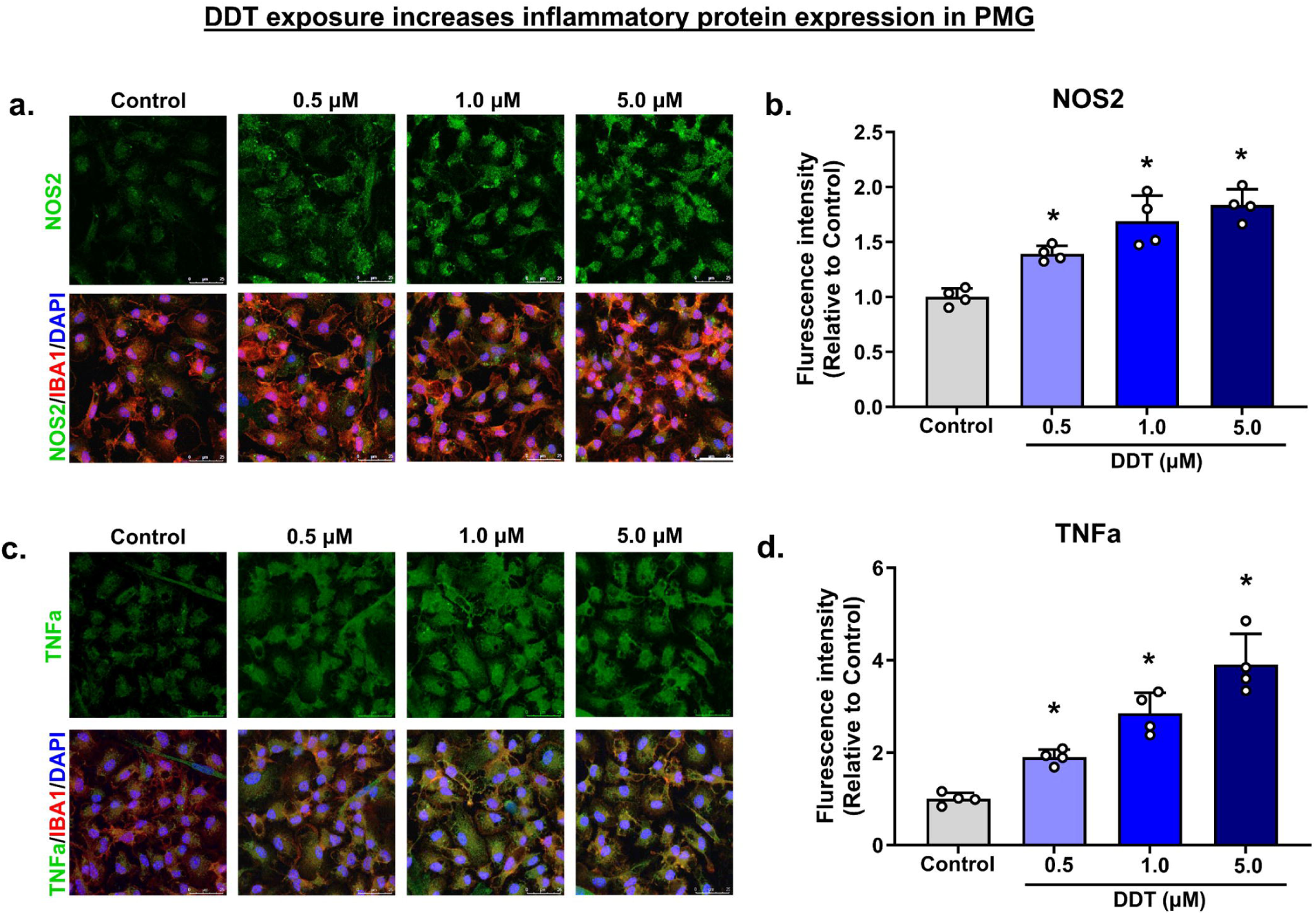
DDT exposure increases NOS2 and TNFa protein levels in PMG. PMG were directly exposed to DDT for 24 h. a) Representative staining for NOS2 in PMG with increasing DDT exposure, b) Quantification of fluorescence intensity/ cell presented as percent control for NOS2. c) Representative staining for TNFa following DDT exposure, d) Quantification of fluorescence intensity/ cell presented as percent control for TNFa. n=4 independent isolations consisting of 3 technical replicates each. Data are presented as means and error bars as ± SD. Statistical significance was determined using a One-way ANOVA. * indicates p<0.05.

### DDT-induced increase in pro-inflammatory cytokines is blocked by TTX

To examine if the increase in pro-inflammatory mRNA involved the VGSCs, we utilized 1 μM tetrodotoxin (TTX) as an antagonist of DDT’s actions as previously described [29]. To assess whether TTX attenuated the pro-inflammatory profile of PMG, cells were treated with 1 μM DDT, as it was the lowest concentration that produced a significant increase in pro-inflammatory genes in the presence or absence of 1 μM TTX. We evaluated this using two-way ANOVA, and examined the main effects of the DDT treatment and TTX **(Figure 4)**. Two-way ANOVA revealed a significant main effect of treatment for *Il-1b* (F_1,12_ =88.76, p<0.0001), *Il-6* (F_1,12_ =6.150, p=0.03), *Nos2* (F_1,12_ =23.07, p=0.0004) and *Tnfa* (F_1,12_ =48.65, p<0.0001); significant main effects of TTX for *Il-1b* (F_1,12_ =28.3, p=0.0002), *Nos2* (F_1,12_ =7.735, p=0.02) and *Tnfa* (F_1,12_ =32.95, p<0.0001); and finally a significant interaction effect for *Il-1b* (F_1,12_ =94.39, p<0.0001), *Il-6* (F_1,12_ =24.07, p=0.0004), *Nos2* (F_1,12_ =22.01, p=0.0005) and *Tnfa* (F_1,12_ =37.02, p<0.0001) **(Figure 4 a-d)**. For all the indicated pro-inflammatory genes upregulated by DDT at 1 μM, pretreatment with TTX abolished this effect. The attenuation of this upregulation of pro-inflammatory genes by TTX demonstrates that the mechanism of action of DDT likely involves modulation of the VGSCs.

**Figure 4:**
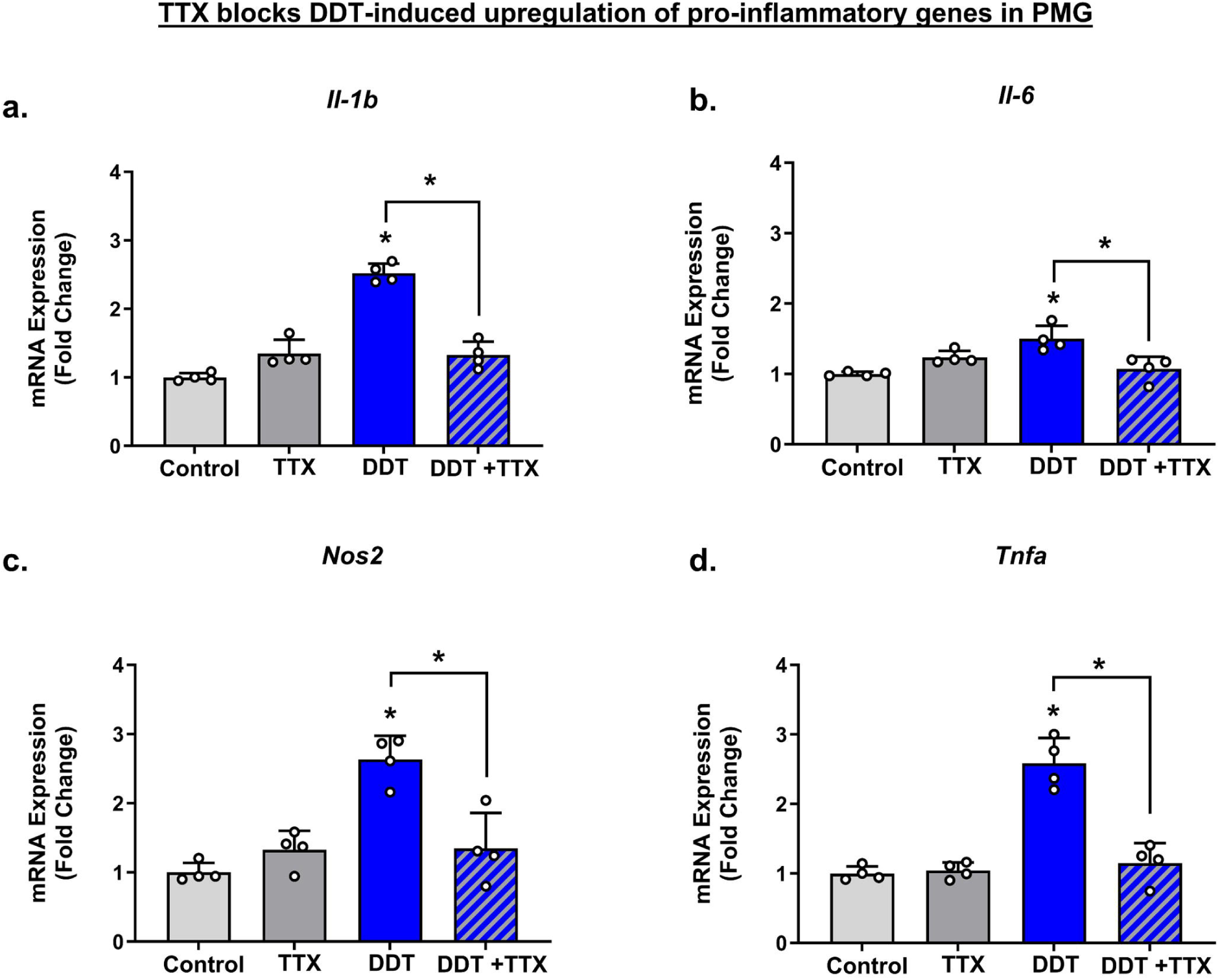
TTX blocks the increase in mRNA expression of proinflammatory genes in PMG. mRNA levels of inflammatory genes in PMG exposed to DDT for 6 h (a - d). n=4 independent isolations consisting of 3 technical replicates each. Data are presented as means and error bars as ± SD. Statistical significance was determined using a Two-way ANOVA. * indicates p<0.05 compared to the control unless indicated otherwise.

### Acute DDT exposure increases the expression of pro-inflammatory genes in vivo

To investigate if exposure to DDT is associated with a neuroinflammatory response *in-vivo,* C57BL/6J male and female mice were exposed to a single dose of DDT (30 mg/kg, oral gavage), and pro-inflammatory gene expression was examined 24 hours later in the hippocampus and frontal cortex. These regions were chosen because of their vulnerability in AD [43], abundant expression of VGSCs [44], and because of our previous work in C57BL6/J mice, which demonstrated changes in the amyloid-β precursor protein in these regions with a sub-chronic lower exposure of DDT (3 mg/kg every 3 days for 1 month) [45]. In the frontal cortex of males, a 1.8-fold increase in *Il-1b* (t=3.993, df=8, p=0.004), 1.5-fold increase in *Il-6* (t=3.042, df=8, p=0.02), 2.3-fold increase in *Nos2* (t=7.360, df=8, p<0.0001), and Tnfa (t=4.292, df=8, p=0.003) **(Figure 5a)** was observed. In the hippocampus, we measured a significant a 1.6-fold increase in *Il-1b* (t=2.982, df=7, p=0.02), 1.4-fold increase in *Il-6* (t=2.729, df=8, p=0.03), 1.7-fold increase in *Nos2* (t=3.698, df=8, p=0.006), and 1.6-fold increase in Tnfa (t=2.841, df=9, p=0.02) **(Figure 5b)**. Similarly, in the female frontal cortex, a 1.5-fold increase in *Il-1b* (t=2.432, df=9, p=0.04), a 1.47-fold increase in *Nos2* (t=2.379, df=8, p=0.04), and 1.6-fold increase in *Tnfa* (t=3.722, df=8, p=0.006) was observed **(Figure 5c)**. Lastly, in the hippocampus of females, we detected an approximately 1.5-fold increase in *Il-1b* (t=3.350, df=9, p=0.0085), a 1.9-fold increase in *Nos2* (t=4.543, df=9, p=0.0014), and a 1.5-fold increase in *Tnfa* (t=3.017, df=9, p=0.0145) **(Figure 5d)**. These data suggest that an acute one-time exposure to DDT can upregulate pro-inflammatory genes in these brain regions.

**Figure 5:**
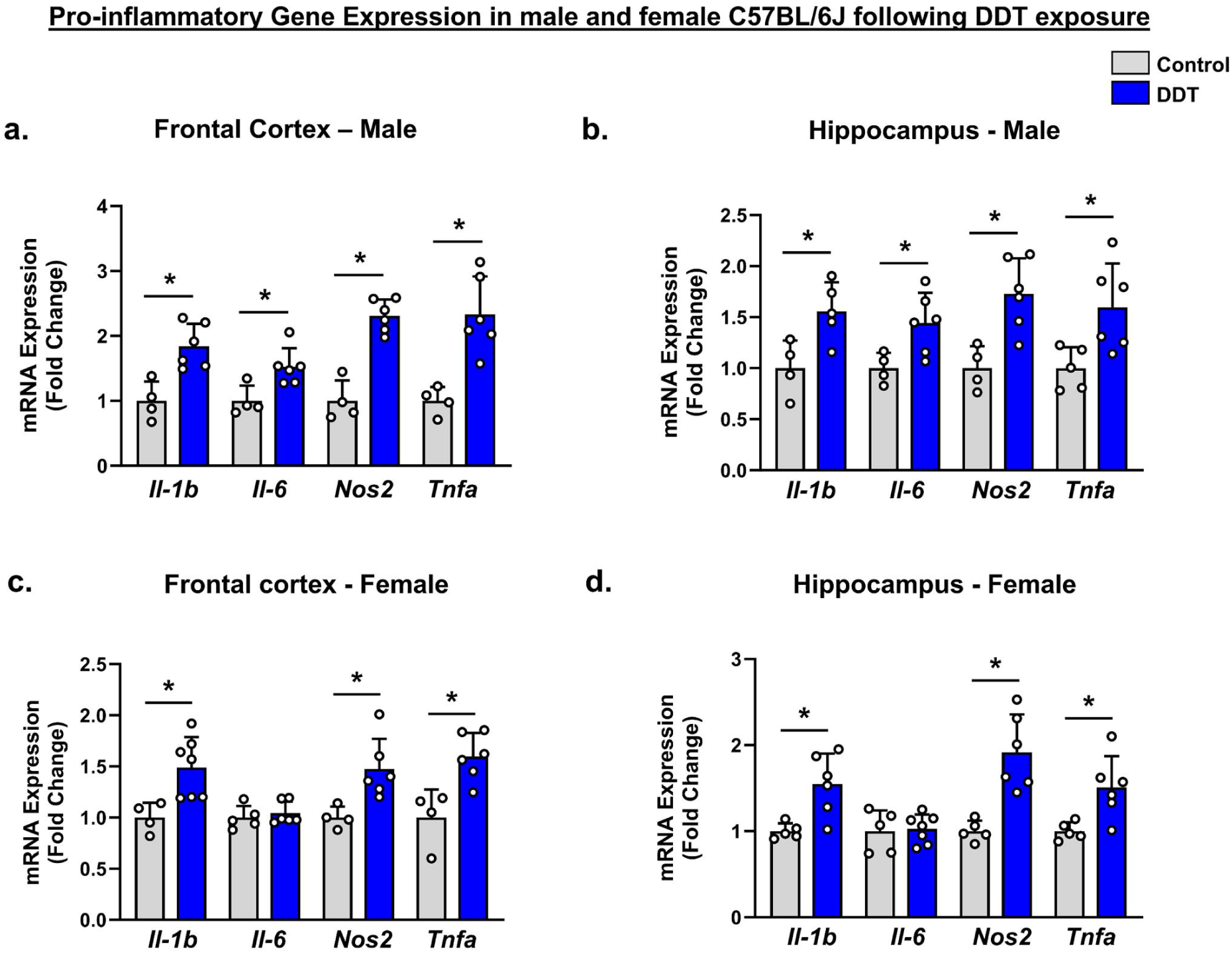
Acute DDT exposure increases proinflammatory gene expression. C57BL/6J animals were exposed to vehicle (corn oil) or a single dose of DDT (30 mg/kg) by oral gavage. mRNA levels of inflammatory genes in the frontal cortex and hippocampus of C57BL/6J males (a and b) and females (c - d). n = 4-7. Data are presented as means and error bars as ± SD. Statistical significance between vehicle control and DDT-exposed mice for each gene was determined using Student’s t-test. * indicates p<0.05.

### DDT exposure alters DAM genes in PMG and in vivo

We first focused on the expression of DAM genes in primary microglia exposed to incremental concentrations of DDT. A significant treatment effect was observed for homeostatic genes, including *Cx3cr1* (F_3,12_ =6.555, p=0.007), *P2ry12* (F_3,12_ =18.98, p<0.0001) and *Tmem119* (F_3,12_ =7.241, p=0.005) **(Figure 6a)**. 1 µM DDT downregulated *Cx3cr1* by 0.8-fold (p=0.02), *P2ry12* by 0.9-fold (p=0.009) and *Tmem119* by 0.8-fold (p=0.02). PMG treated with 5 µM DDT, further decreased the expression of *Cx3cr1* by 0.7-fold (p=0.003), *P2ry12* by 0.8-fold (p<0.0001) and *Tmem119* by 0.7-fold (p=0.002). Subsequent analysis of stage 1 DAM genes that are Trem2-independent, revealed a significant treatment effect for *Ctsb* (F_3,12_ =26.11, p<0.0001), *Tyrobp* (F_3,12_ =24.25, p<0.0001) and *Apoe* (F_3,12_ =65.99, p<0.0001) **(Figure 6b)**. PMG treated with 0.5, 1 and 5 µM DDT, showed a steady concentration-dependent increase in mRNA levels. *Ctsb* increased 1.15-fold (0.5 µM; p=0.003), 1.21-fold (1 µM; p=0.0002) and 1.32-fold (5 µM; p<0.0001). *Tyrobp* levels increased by 1.16-fold (0.5 µM; p=0.009), 1.26-fold (1 µM; p=0.0002) and 1.36-fold (5 µM; p<0.0001). Lastly, *Apoe* mRNA levels significantly increased by 1.43-fold (0.5 µM; p=0.008), 2.01-fold (1 µM; p<0.0001) and 2.56-fold (5 µM; p<0.0001). Next, we measured mRNA levels of *Trem2*, in order to assess changes in Trem2-dependent DAM genes. A significant DDT treatment effect was observed (F_3,12_ =12.96, p=0.0005), with statistically significant 1.2-fold upregulation measured only at 5 µM (p=0.004).

**Figure 6:**
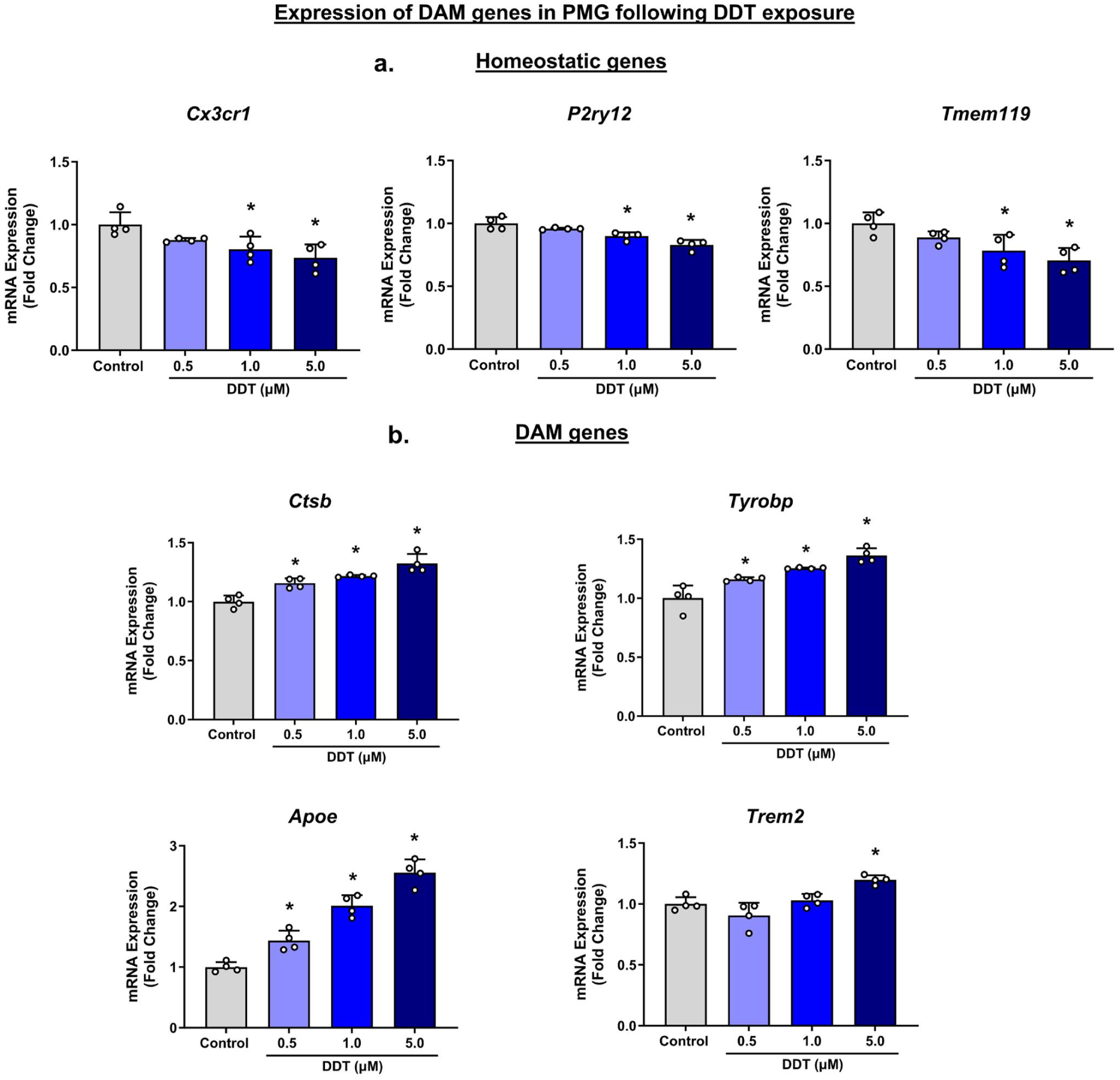
DDT exposure alters gene expression of disease-associated microglia. mRNA levels of homeostatic genes (a) and disease-associated microglia (DAM) genes (b) in PMG exposed to DDT for 6h. n=4 independent isolations consisting of 3 technical replicates each. Data are presented as means and error bars as ± SD. Statistical significance was determined using a One-way ANOVA. * indicates p<0.05.

We further evaluated the expression of DAM genes in the frontal cortex and hippocampus of male and female mice acutely exposed to DDT. In the frontal cortex of males, a 0.9-fold decrease in *P2ry12* (t=2.516, df=10, p=0.03), 0.8-fold decrease in *Tmem119* (t=3.338, df=9, p=0.009), and an approximate 1.3-fold increase in *Ctsb* (t=4.309, df=10, p=0.002) and *Tyrobp* (t=6.047, df=10, p=0.0001) **(Figure 7a)** was observed. In the hippocampus we measured a significant decrease of 0.8-fold in *Cx3cr1* (t=3.880, df=9, p=0.004), 0.9-fold in *P2ry12* (t=2.363, df=10, p=0.04), and 0.8-fold in *Tmem119* (t=3.588, df=10, p=0.005), whereas a 1.4-fold increase in *Ctsb* (t=3.511, df=9, p=0.007) and 1.2-fold increase in *Tyrobp* (t=2.922, df=9, p=0.02) **(Figure 7b)**. No changes were noted in the expression of *Apoe* and *Trem2*. In the female frontal cortex, we observed a significant decrease of 0.7-fold in *Cx3cr1* (t=4.898, df=11, p=0.0005), 0.9-fold in *P2ry12* (t=4.919, df=11, p=0.0005), and 0.8-fold in *Tmem119* (t=3.457, df=11, p=0.005). Conversely, a significant increase was observed in the expression of *Ctsb* by 1.4-fold (t=8.244, df=11, p<0.0001), *Tyrobp* by 1.2-fold (t=5.531, df=12, p=0.0001), *Apoe* by 1.3-fold (t=4.172, df=12, p=0.001) and *Trem2* by 1.3-fold (t=3.808, df=11, p=0.003) **(Figure 7c)**. Lastly, in the hippocampus of females, we detected an approximate 0.9-fold decrease in *Cx3cr1* (t=2.312, df=11, p=0.04), *P2ry12* (t=4.932, df=12, p=0.0003), and *Tmem119* (t=4.148, df=12, p=0.001), whereas a significant increase of 1.3-fold in *Ctsb* (t=3.733, df=11, p=0.003), 1.2-fold in *Tyrobp* (t=6.373, df=12, p<0.0001), 1.6-fold in *Apoe* (t=5.908, df=10, p=0.0002) and 1.2-fold in *Trem2* was observed **(Figure 7d)**.

**Figure 7:**
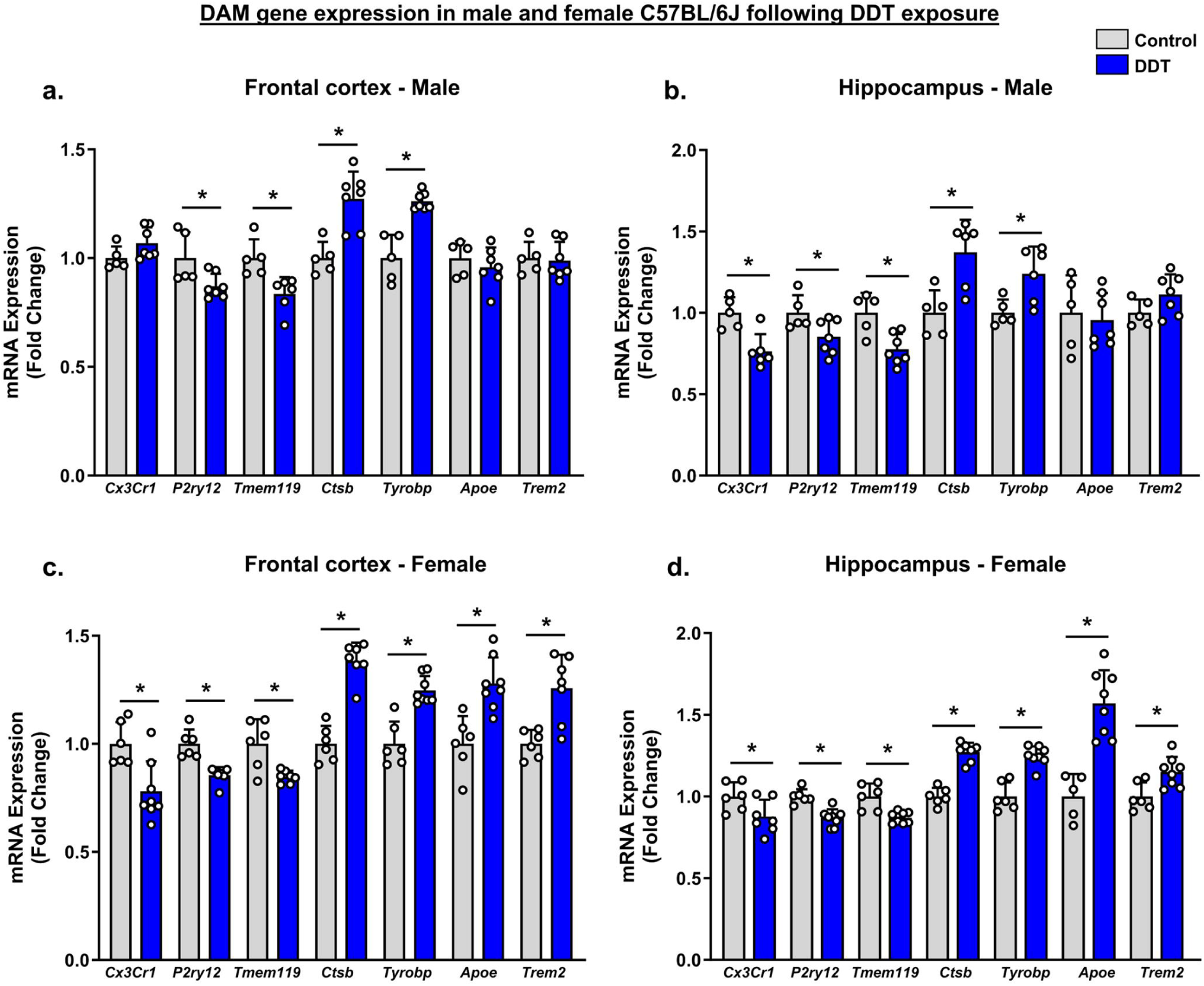
Acute DDT exposure alters levels of DAM genes. C57BL/6J animals were exposed to vehicle (corn oil) or a single dose of DDT (30 mg/kg) by oral gavage. mRNA levels of DAM genes in the frontal cortex and hippocampus of C57BL/6J males (a and b) and females (c - d). n = 5-8. Data are presented as means and error bars as ± SD. Statistical significance between vehicle control and DDT-exposed mice for each gene was determined using Student’s t-test. * indicates p<0.05.

## Discussion

Neurodegenerative diseases such as Alzheimer’s are complex, with many pathways contributing to disease progression and pathologies [46, 47]. These diseases typically involve the aggregation of pathological proteins, degeneration of neurons, and chronic neuroinflammation [48, 49]. Studies on post-mortem brains, CSF levels and blood from AD patients have demonstrated that neuroinflammation in AD is not only an effect of pathogenesis but is closely involved in the progression and accumulation of senile plaques and neurofibrillary tangles [50-53]. Neuroinflammation has emerged as a prominent player in AD research, with the identification of more than 20 gene variants associated with AD by recent GWAS studies [54]. Several of these genes, such as *TREM2, CR1, MS4A, CD33* and *CLU*, regulate immune and inflammatory responses, and some are directly associated with microglia [55, 56]. Recent evidence suggests that microglia and astrocytes, may become activated by environmental stressors, including but not limited to metals, pesticides, and components of air pollutants [57-59].

Sporadic Alzheimer’s disease is a complex and multi-faceted disease with significant contributions from environmental factors, including toxicant exposures [60]. Recent epidemiological evidence suggests that DDT and its metabolite dichlorodiphenyldichloroethylene (DDE) may increase the risk of AD [61-64]. Previously, we reported four-fold higher serum levels of DDE in AD patients, with a corresponding increased risk of AD (Odds ratio=4.18) [63]. While the mechanisms behind this risk remain under investigation, our laboratory has recently reported that DDT can upregulate critical genes involved in the amyloidogenic pathway in multiple experimental models and accelerate AD-related pathology in the 3xTG-AD mouse model [45]. While there is currently no published evidence for DDT exposure and neuroinflammation, there have been investigations of its inflammatory effects in the periphery. Previous studies by other groups have investigated the effects of DDT on the induction or activation of cells in the periphery, notably by increasing synthesis and or secretion of pro-inflammatory cytokines. DDT exposure (0.025-2.5 uM) increased IL-1b secretion in human peripheral blood mononuclear cells (PBMCs), monocyte-depleted PBMCs (MD-PBMCs) and natural killer (NK) cells [65], and mRNA levels of *IL-1b* in PBMCs [66]. Similarly, DDT exposure increases levels of IFNg in NK, MD-PBMCs and PBMCs, and TNFa secretion from MD-PBMCs but not in NK or PBMCs (0.025-2.5 uM) [67]. Moreover, another study documented similar alterations in isolated PBMCs which were accompanied by glial activation, increased NFκB signaling, and elevated protein levels of TNFa, IFNg, IL-6, and IL-1b [67, 68].

We have previously demonstrated that the pyrethroid insecticides deltamethrin and permethrin directly activate PMG isolated from C57BL6/J mice through sodium influx and increase the production and secretion of pro-inflammatory cytokines, which is attenuated by TTX [29]. PMG express functional VGSC isoforms Nav1.1, Nav1.2, and Nav1.6 that are sensitive to TTX [23, 24, 29]. Other studies have also indicated the expression of TTX-resistant VGSC Nav 1.5 in microglia [19, 21]. Narahashi and colleagues have comprehensively elucidated the mechanism of DDT, detailing its direct inactivation of the transition of Nav channels into the closed and inactivated states within neurons [69-71]. Because of this shared mechanism of action and findings that both classes of pesticides can act on AD pathways, we sought to examine the effects of DDT on neuroinflammation.

Here, we investigated the effects of DDT on gene expression of pro-inflammatory factors in isolated glial cells and in vivo in wild-type mice. DDT was extensively used in the United States from the 1940s to the 1970s until it was banned in the US in 1972 [26, 27]. Because of its potent actions, it is still used in several developing countries. As an insecticide, its primary uses are both for agricultural applications as well as to aid in public health concerns as it has been highly efficacious against malaria outbreaks [26]. Both DDT and its stable metabolite dichlorodiphenyldichloroethylene (DDE) have a long half-life, high lipophilicity, and low reactivity [27, 28]. These properties allow DDT and its metabolites to persist over a long period in the environment. This, coupled with the slow elimination of DDT from organisms, allows for bioaccumulation [28]. A recent US Food and Drug Administration report found measurable levels of DDT and its metabolites in 2.2 % of surveyed domestic food samples and 0.6 % of imported food samples [72]. Furthermore, according to the Center for Disease Control and Prevention’s National Health Survey, levels of DDE are still found in human serum samples [73, 74]. Furthermore, while DDE levels have rapidly declined in many countries, it is still classified as high in human samples in specific geographic regions, including Eastern Europe, Central America, and Eastern Asia, suggesting it is still a paramount environmental health concern [75].

The concentrations of DDT used for the *in vitro* studies were the same as in our previous publications [45]. Here, we demonstrated that DDT acts on microglia to upregulate pro-inflammatory factors, including *Il-1b*, *Tnfa,* and *Nos2*. Furthermore, we provide evidence that this may be specific to microglia, as no significant increase in pro-inflammatory genes was measured when PMA were treated with DDT. A well-established and known mechanism of DDT is persistent depolarization via modulation of the VGSCs [76]. To examine if the ability of DDT to increase pro-inflammatory factors is related to its modulation of the VGSC, we pre-treated microglial cells with the VGSC antagonist TTX as previously described [45]. The dose for TTX (1μM) was chosen to block TTX-sensitive VGSCs and has been previously used in our laboratory on PMG [23, 29, 77]. The inclusion of TTX abolished the increased gene expression of pro-inflammatory factors observed with DDT alone, suggesting that this direct action likely involves the VGSC. This is consistent with previously published work with pyrethroids [29]. Previous studies have documented the predominant expression of TTX-resistant VGSCs, particularly Nav1.5 in astrocytes [20]. However, reports have also indicated the expression of TTX-sensitive VGSCs Nav1.2, Nav1.3, and Nav1.6, albeit at lower levels [78, 79]. This could potentially explain the lack of direct effects of DDT on astrocytes. Next, we assessed pro-inflammatory gene expression in the hippocampus and frontal cortex, two particularly vulnerable regions in AD. Similar to PMG, DDT exposure increased the mRNA expression of *Il-1b, Nos2* and *Tnfa* in the hippocampus and frontal cortex of both males and females. However, post DDT exposure, mRNA levels of *Il-6* increased only in the hippocampus and frontal cortex of males but not females. This finding is significant because of the relatively short exposure time where animals were sacrificed at 24 hours. These data provide the first evidence that acute exposure in wild-type mice causes upregulation of pro-inflammatory pathways. Rodent male and female microglia differ in numbers, response to external stimuli and other functions including phagocytosis, synaptic pruning across development, age, and brain regions [80]. Thus the difference in proinflammatory response across brain regions and sex could be attributed to the age at which the mice were exposed.

Evidence from previous studies has shown that transcriptional profiles of microglia shift from homeostatic phenotype to inflammatory profiles, including disease-associated microglia (DAM). This microglial phenotype is observed in aging [81] and neurodegenerative diseases, such as AD and ALS [15, 81], as well as exposure to air particulate matter (PM 2.5) [82] and diesel exhaust [83]. Importantly, DAM are characterized by the downregulation of homeostatic genes along with an upregulation of genes that are involved in lipid metabolism, and signaling pathways for inflammation and phagocytosis [81]. The transition of homeostatic microglia to DAM is a sequential process, where the first shift from homeostasis to stage 1 DAM is Trem2-independent, while subsequent conversion of stage 1 DAM to stage 2 DAM is dependent on Trem2 signaling [15, 16]. We observed a uniform downregulation in the gene expression of homeostatic genes, including *Cx3cr1*, *P2ry12* and *Tmem119* in PMG exposed to DDT and a concentration-dependent upregulation of stage 1 DAM genes, including *Ctsb*, *Tyrobp* and *Apoe*. Upregulation in *Trem2* was only observed at 5 µM DDT. In the frontal cortex and hippocampus of male and female mice acutely exposed to DDT, we observed a downregulation of homeostatic genes and an upregulation of *Ctsb* and *Tyrobp*. Interestingly, *Apoe* and *Trem2* mRNA levels only increased in the brain regions of females exposed to DDT. This suggests that – (1) DDT activates stage 1 DAM genes in both males and females, (2) Only females acutely exposed to DDT, showed increased expression of *Apoe* and *Trem2* that indicates potential sex-related regulation of DAM genes and Trem2 regulators, *Tyrobp* and *Apoe*, following DDT exposure. Recent evidence in aged wildtype and AD mouse models has shown that specific DAM gene expression was altered only in female mice [84, 85]. Other studies have shown that homeostatic microglia switch to activated response microglia (ARMs) more rapidly in females compared to males [86]. In post-mortem AD human brain tissue, a morphological difference was observed between male and female microglia, with lower microglial density in female tissue compared to males [87]. In another study, Mathys and colleagues showed that a subpopulation of AD-associated microglia were enriched in samples from females [88]. Evidence in humans also indicates microglial genes including TREM2 showed profound dysregulation in females compared to males in the temporal cortex, parahippocampal gyrus, and cerebellum of human AD tissue [89].

The data reported here were generated in wild-type C57BL/6J mice, further study is needed in transgenic models that produce plaque and/or tangle pathology. Moreover, the effect of DDT on neuronal VGSCs can indirectly influence microglial activity through alterations in excitotoxicity, extracellular ATP levels, and synaptic activity [24, 90]. Additional studies are required to explore indirect mechanisms of neuronal activity on microglial regulation post-DDT exposure. To our knowledge, this is the first study evaluating the novel hypothesis that DDT can directly activate microglia and investigating neuroinflammation in an animal model following DDT exposure and. These data demonstrate that exposure to DDT is associated with an increased pro-inflammatory response and enhanced DAM profile *in vivo* and *in vitro*. Furthermore, these results suggest that this effect is microglial specific, and DDT does not directly activate astrocytes. Lastly, we also show evidence that VGSCs modulate this process, at least *in vitro*, as the effect of DDT was blocked by the inclusion of TTX. These findings, coupled with our previous data examining the effect of DDT on the amyloidogenic pathway, demonstrate that DDT likely acts on multiple mechanisms involved in AD, including neuroinflammation.

## Author Contributions

Isha Mhatre-Winters (Conceptualization; Methodology; Formal analysis; Data curation; Writing - Original Draft); Aseel Eid (Conceptualization; Methodology; Formal analysis; Writing - Original Draft); Nicole Blum (Methodology; Formal analysis); Yoonhee Han (Methodology); Ferass M. Sammoura (Methodology); Long-Jun Wu (Writing-Review and Editing; Funding acquisition) and Jason R. Richardson (Conceptualization; Formal analysis; Writing - Review & Editing; Funding acquisition). All authors have read and agreed to the published version of the manuscript.

## Acknowledgments

The authors have no acknowledgments to report.

## Datasets/ Data Availability Statement

The data supporting the findings of this study are available within the article and/or its supplementary material.

## References

[1] Lambert JC, Ibrahim-Verbaas CA, Harold D, Naj AC, Sims R, Bellenguez C, DeStafano AL, Bis JC, Beecham GW, Grenier-Boley B, Russo G, Thorton-Wells TA, Jones N, Smith AV, Chouraki V, Thomas C, Ikram MA, Zelenika D, Vardarajan BN, Kamatani Y, Lin CF, Gerrish A, Schmidt H, Kunkle B, Dunstan ML, Ruiz A, Bihoreau MT, Choi SH, Reitz C, Pasquier F, Cruchaga C, Craig D, Amin N, Berr C, Lopez OL, De Jager PL, Deramecourt V, Johnston JA, Evans D, Lovestone S, Letenneur L, Moron FJ, Rubinsztein DC, Eiriksdottir G, Sleegers K, Goate AM, Fievet N, Huentelman MW, Gill M, Brown K, Kamboh MI, Keller L, Barberger-Gateau P, McGuiness B, Larson EB, Green R, Myers AJ, Dufouil C, Todd S, Wallon D, Love S, Rogaeva E, Gallacher J, St George-Hyslop P, Clarimon J, Lleo A, Bayer A, Tsuang DW, Yu L, Tsolaki M, Bossu P, Spalletta G, Proitsi P, Collinge J, Sorbi S, Sanchez-Garcia F, Fox NC, Hardy J, Deniz Naranjo MC, Bosco P, Clarke R, Brayne C, Galimberti D, Mancuso M, Matthews F, European Alzheimer’s Disease I, Genetic, Environmental Risk in Alzheimer’s D, Alzheimer’s Disease Genetic C, Cohorts for H, Aging Research in Genomic E, Moebus S, Mecocci P, Del Zompo M, Maier W, Hampel H, Pilotto A, Bullido M, Panza F, Caffarra P, Nacmias B, Gilbert JR, Mayhaus M, Lannefelt L, Hakonarson H, Pichler S, Carrasquillo MM, Ingelsson M, Beekly D, Alvarez V, Zou F, Valladares O, Younkin SG, Coto E, Hamilton-Nelson KL, Gu W, Razquin C, Pastor P, Mateo I, Owen MJ, Faber KM, Jonsson PV, Combarros O, O’Donovan MC, Cantwell LB, Soininen H, Blacker D, Mead S, Mosley TH, Jr., Bennett DA, Harris TB, Fratiglioni L, Holmes C, de Bruijn RF, Passmore P, Montine TJ, Bettens K, Rotter JI, Brice A, Morgan K, Foroud TM, Kukull WA, Hannequin D, Powell JF, Nalls MA, Ritchie K, Lunetta KL, Kauwe JS, Boerwinkle E, Riemenschneider M, Boada M, Hiltuenen M, Martin ER, Schmidt R, Rujescu D, Wang LS, Dartigues JF, Mayeux R, Tzourio C, Hofman A, Nothen MM, Graff C, Psaty BM, Jones L, Haines JL, Holmans PA, Lathrop M, Pericak-Vance MA, Launer LJ, Farrer LA, van Duijn CM, Van Broeckhoven C, Moskvina V, Seshadri S, Williams J, Schellenberg GD, Amouyel P (2013) Meta-analysis of 74,046 individuals identifies 11 new susceptibility loci for Alzheimer’s disease. Nat Genet 45, 1452-1458.

[2] Li JQ, Wang HF, Zhu XC, Sun FR, Tan MS, Tan CC, Jiang T, Tan L, Yu JT, Alzheimer’s Disease Neuroimaging I (2017) GWAS-Linked Loci and Neuroimaging Measures in Alzheimer’s Disease. Mol Neurobiol 54, 146–153.

[3] Zhang X, Zhu C, Beecham G, Vardarajan BN, Ma Y, Lancour D, Farrell JJ, Chung J, Alzheimer’s Disease Sequencing P, Mayeux R, Haines JL, Schellenberg GD, Pericak-Vance MA, Lunetta KL, Farrer LA (2019) A rare missense variant of CASP7 is associated with familial late-onset Alzheimer’s disease. Alzheimers Dement 15, 441-452.

[4] Patel D, Mez J, Vardarajan BN, Staley L, Chung J, Zhang X, Farrell JJ, Rynkiewicz MJ, Cannon-Albright LA, Teerlink CC, Stevens J, Corcoran C, Gonzalez Murcia JD, Lopez OL, Mayeux R, Haines JL, Pericak-Vance MA, Schellenberg G, Kauwe JSK, Lunetta KL, Farrer LA, Alzheimer’s Disease Sequencing P (2019) Association of Rare Coding Mutations With Alzheimer Disease and Other Dementias Among Adults of European Ancestry. JAMA Netw Open 2, e191350.

[5] Ma Y, Jun GR, Zhang X, Chung J, Naj AC, Chen Y, Bellenguez C, Hamilton-Nelson K, Martin ER, Kunkle BW, Bis JC, Debette S, DeStefano AL, Fornage M, Nicolas G, van Duijn C, Bennett DA, De Jager PL, Mayeux R, Haines JL, Pericak-Vance MA, Seshadri S, Lambert JC, Schellenberg GD, Lunetta KL, Farrer LA, Alzheimer’s Disease Sequencing P, Alzheimer’s Disease Exome Sequencing-France P (2019) Analysis of Whole-Exome Sequencing Data for Alzheimer Disease Stratified by APOE Genotype. JAMA Neurol.

[6] Bis JC, Jian X, Kunkle BW, Chen Y, Hamilton-Nelson KL, Bush WS, Salerno WJ, Lancour D, Ma Y, Renton AE, Marcora E, Farrell JJ, Zhao Y, Qu L, Ahmad S, Amin N, Amouyel P, Beecham GW, Below JE, Campion D, Charbonnier C, Chung J, Crane PK, Cruchaga C, Cupples LA, Dartigues JF, Debette S, Deleuze JF, Fulton L, Gabriel SB, Genin E, Gibbs RA, Goate A, Grenier-Boley B, Gupta N, Haines JL, Havulinna AS, Helisalmi S, Hiltunen M, Howrigan DP, Ikram MA, Kaprio J, Konrad J, Kuzma A, Lander ES, Lathrop M, Lehtimaki T, Lin H, Mattila K, Mayeux R, Muzny DM, Nasser W, Neale B, Nho K, Nicolas G, Patel D, Pericak-Vance MA, Perola M, Psaty BM, Quenez O, Rajabli F, Redon R, Reitz C, Remes AM, Salomaa V, Sarnowski C, Schmidt H, Schmidt M, Schmidt R, Soininen H, Thornton TA, Tosto G, Tzourio C, van der Lee SJ, van Duijn CM, Vardarajan B, Wang W, Wijsman E, Wilson RK, Witten D, Worley KC, Zhang X, Alzheimer’s Disease Sequencing P, Bellenguez C, Lambert JC, Kurki MI, Palotie A, Daly M, Boerwinkle E, Lunetta KL, Destefano AL, Dupuis J, Martin ER, Schellenberg GD, Seshadri S, Naj AC, Fornage M, Farrer LA (2018) Whole exome sequencing study identifies novel rare and common Alzheimer’s-Associated variants involved in immune response and transcriptional regulation. Mol Psychiatry.

[7] Serrano-Pozo A, Aldridge GM, Zhang Q (2017) Four Decades of Research in Alzheimer’s Disease (1975-2014): A Bibliometric and Scientometric Analysis. J Alzheimers Dis 59, 763–783.

[8] Serrano-Pozo A, Mielke ML, Gomez-Isla T, Betensky RA, Growdon JH, Frosch MP, Hyman BT (2011) Reactive glia not only associates with plaques but also parallels tangles in Alzheimer’s disease. Am J Pathol 179, 1373–1384.

[9] Spangenberg EE, Green KN (2017) Inflammation in Alzheimer’s disease: Lessons learned from microglia-depletion models. Brain Behav Immun 61, 1–11.

[10] Domingues C, da Cruz ESOAB, Henriques AG (2017) Impact of Cytokines and Chemokines on Alzheimer’s Disease Neuropathological Hallmarks. Curr Alzheimer Res 14, 870-882.

[11] Tang Y, Le W (2016) Differential Roles of M1 and M2 Microglia in Neurodegenerative Diseases. Mol Neurobiol 53, 1181–1194.

[12] Ransohoff RM (2016) A polarizing question: do M1 and M2 microglia exist? Nat Neurosci 19, 987–991.

[13] Paolicelli RC, Sierra A, Stevens B, Tremblay ME, Aguzzi A, Ajami B, Amit I, Audinat E, Bechmann I, Bennett M, Bennett F, Bessis A, Biber K, Bilbo S, Blurton-Jones M, Boddeke E, Brites D, Brone B, Brown GC, Butovsky O, Carson MJ, Castellano B, Colonna M, Cowley SA, Cunningham C, Davalos D, De Jager PL, de Strooper B, Denes A, Eggen BJL, Eyo U, Galea E, Garel S, Ginhoux F, Glass CK, Gokce O, Gomez-Nicola D, Gonzalez B, Gordon S, Graeber MB, Greenhalgh AD, Gressens P, Greter M, Gutmann DH, Haass C, Heneka MT, Heppner FL, Hong S, Hume DA, Jung S, Kettenmann H, Kipnis J, Koyama R, Lemke G, Lynch M, Majewska A, Malcangio M, Malm T, Mancuso R, Masuda T, Matteoli M, McColl BW, Miron VE, Molofsky AV, Monje M, Mracsko E, Nadjar A, Neher JJ, Neniskyte U, Neumann H, Noda M, Peng B, Peri F, Perry VH, Popovich PG, Pridans C, Priller J, Prinz M, Ragozzino D, Ransohoff RM, Salter MW, Schaefer A, Schafer DP, Schwartz M, Simons M, Smith CJ, Streit WJ, Tay TL, Tsai LH, Verkhratsky A, von Bernhardi R, Wake H, Wittamer V, Wolf SA, Wu LJ, Wyss-Coray T (2022) Microglia states and nomenclature: A field at its crossroads. Neuron 110, 3458–3483.

[14] Wang J, He W, Zhang J (2023) A richer and more diverse future for microglia phenotypes. Heliyon 9, e14713.

[15] Keren-Shaul H, Spinrad A, Weiner A, Matcovitch-Natan O, Dvir-Szternfeld R, Ulland TK, David E, Baruch K, Lara-Astaiso D, Toth B, Itzkovitz S, Colonna M, Schwartz M, Amit I (2017) A Unique Microglia Type Associated with Restricting Development of Alzheimer’s Disease. Cell 169, 1276–1290 e1217.

[16] Deczkowska A, Keren-Shaul H, Weiner A, Colonna M, Schwartz M, Amit I (2018) Disease-Associated Microglia: A Universal Immune Sensor of Neurodegeneration. Cell 173, 1073–1081.

[17] Butovsky O, Jedrychowski MP, Moore CS, Cialic R, Lanser AJ, Gabriely G, Koeglsperger T, Dake B, Wu PM, Doykan CE, Fanek Z, Liu L, Chen Z, Rothstein JD, Ransohoff RM, Gygi SP, Antel JP, Weiner HL (2014) Identification of a unique TGF-beta-dependent molecular and functional signature in microglia. Nat Neurosci 17, 131–143.

[18] Friedman BA, Srinivasan K, Ayalon G, Meilandt WJ, Lin H, Huntley MA, Cao Y, Lee SH, Haddick PCG, Ngu H, Modrusan Z, Larson JL, Kaminker JS, van der Brug MP, Hansen DV (2018) Diverse Brain Myeloid Expression Profiles Reveal Distinct Microglial Activation States and Aspects of Alzheimer’s Disease Not Evident in Mouse Models. Cell Rep 22, 832–847.

[19] Black JA, Liu S, Waxman SG (2009) Sodium channel activity modulates multiple functions in microglia. Glia 57, 1072–1081.

[20] Pappalardo LW, Black JA, Waxman SG (2016) Sodium channels in astroglia and microglia. Glia 64, 1628–1645.

[21] Black JA, Waxman SG (2012) Sodium channels and microglial function. Exp Neurol 234, 302–315.

[22] Richardson JR, Hossain MM (2013) Microglial ion channels as potential targets for neuroprotection in Parkinson’s disease. Neural Plast 2013, 587418.

[23] Hossain MM, Sonsalla PK, Richardson JR (2013) Coordinated role of voltage-gated sodium channels and the Na+/H+ exchanger in sustaining microglial activation during inflammation. Toxicol Appl Pharmacol 273, 355–364.

[24] Jung GY, Lee JY, Rhim H, Oh TH, Yune TY (2013) An increase in voltage-gated sodium channel current elicits microglial activation followed inflammatory responses in vitro and in vivo after spinal cord injury. Glia 61, 1807–1821.

[25] Soderlund DM (2012) Molecular mechanisms of pyrethroid insecticide neurotoxicity: recent advances. Arch Toxicol 86, 165–181.

[26] Bate R (2007) The Rise, Fall, Rise, and Imminent Fall of DDT. American Enterprise Institute for Public Policy Research 14, 1–9.

[27] Turusov V, Rakitsky V, Tomatis L (2002) Dichlorodiphenyltrichloroethane (DDT): ubiquity, persistence, and risks. Environ Health Perspect 110, 125–128.

[28] Beard J, Australian Rural Health Research C (2006) DDT and human health. Sci Total Environ 355, 78–89.

[29] Hossain MM, Liu J, Richardson JR (2017) Pyrethroid Insecticides Directly Activate Microglia Through Interaction With Voltage-Gated Sodium Channels. Toxicol Sci 155, 112–123.

[30] Percie du Sert N, Ahluwalia A, Alam S, Avey MT, Baker M, Browne WJ, Clark A, Cuthill IC, Dirnagl U, Emerson M, Garner P, Holgate ST, Howells DW, Hurst V, Karp NA, Lazic SE, Lidster K, MacCallum CJ, Macleod M, Pearl EJ, Petersen OH, Rawle F, Reynolds P, Rooney K, Sena ES, Silberberg SD, Steckler T, Wurbel H (2020) Reporting animal research: Explanation and elaboration for the ARRIVE guidelines 2.0. PLoS Biol 18, e3000411.

[31] Council NR (2011) Guide for the Care and Use of Laboratory Animals The National Academic Press, Washington (DC).

[32] Mhatre-Winters I, Eid A, Han Y, Tieu K, Richardson JR (2022) Sex and APOE Genotype Alter the Basal and Induced Inflammatory States of Primary Microglia from APOE Targeted Replacement Mice. Int J Mol Sci 23.

[33] Mhatre-Winters I, Eid A, Han Y, Tieu K, Richardson JR (2023) Sex and APOE Genotype Alter the Basal and Induced Inflammatory States of Primary Astrocytes from Humanized Targeted Replacement Mice. ASN Neuro 15, 17590914221144549.

[34] (EPA) AfTSaDRAaEPA (2022).

[35] Registry A-AfTSaD (2022) Toxicological Profile for DDT, DDE, DDD. U.S. Department of Health and Human Services, Public Health Service Atlanta, GA.

[36] Schmittgen TD, Livak KJ (2008) Analyzing real-time PCR data by the comparative C(T) method. Nat Protoc 3, 1101–1108.

[37] Hu F, Wang Q, Wang P, Wang W, Qian W, Xiao H, Wang L (2012) 17beta-Estradiol regulates the gene expression of voltage-gated sodium channels: role of estrogen receptor alpha and estrogen receptor beta. Endocrine 41, 274–280.

[38] Kow LM, Pfaff DW (2016) Rapid estrogen actions on ion channels: A survey in search for mechanisms. Steroids 111, 46–53.

[39] Dubbelaar ML, Kracht L, Eggen BJL, Boddeke E (2018) The Kaleidoscope of Microglial Phenotypes. Front Immunol 9, 1753.

[40] Cunningham C, Dunne A, Lopez-Rodriguez AB (2019) Astrocytes: Heterogeneous and Dynamic Phenotypes in Neurodegeneration and Innate Immunity. Neuroscientist 25, 455–474.

[41] Kwon HS, Koh SH (2020) Neuroinflammation in neurodegenerative disorders: the roles of microglia and astrocytes. Transl Neurodegener 9, 42.

[42] Garland EF, Hartnell IJ, Boche D (2022) Microglia and Astrocyte Function and Communication: What Do We Know in Humans? Front Neurosci 16, 824888.

[43] Fu H, Hardy J, Duff KE (2018) Selective vulnerability in neurodegenerative diseases. Nat Neurosci 21, 1350–1358.

[44] Whitaker WR, Clare JJ, Powell AJ, Chen YH, Faull RL, Emson PC (2000) Distribution of voltage-gated sodium channel alpha-subunit and beta-subunit mRNAs in human hippocampal formation, cortex, and cerebellum. J Comp Neurol 422, 123–139.

[45] Eid A, Mhatre-Winters I, Sammoura FM, Edler MK, von Stein R, Hossain MM, Han Y, Lisci M, Carney K, Konsolaki M, Hart RP, Bennett JW, Richardson JR (2022) Effects of DDT on Amyloid Precursor Protein Levels and Amyloid Beta Pathology: Mechanistic Links to Alzheimer’s Disease Risk. Environ Health Perspect 130, 87005.

[46] Guo T, Zhang D, Zeng Y, Huang TY, Xu H, Zhao Y (2020) Molecular and cellular mechanisms underlying the pathogenesis of Alzheimer’s disease. Mol Neurodegener 15, 40.

[47] Calabro M, Rinaldi C, Santoro G, Crisafulli C (2021) The biological pathways of Alzheimer disease: a review. AIMS Neurosci 8, 86–132.

[48] Brettschneider J, Del Tredici K, Lee VM, Trojanowski JQ (2015) Spreading of pathology in neurodegenerative diseases: a focus on human studies. Nat Rev Neurosci 16, 109–120.

[49] Guzman-Martinez L, Maccioni RB, Andrade V, Navarrete LP, Pastor MG, Ramos-Escobar N (2019) Neuroinflammation as a Common Feature of Neurodegenerative Disorders. Front Pharmacol 10, 1008.

[50] Kauwe JS, Bailey MH, Ridge PG, Perry R, Wadsworth ME, Hoyt KL, Staley LA, Karch CM, Harari O, Cruchaga C, Ainscough BJ, Bales K, Pickering EH, Bertelsen S, Alzheimer’s Disease Neuroimaging I, Fagan AM, Holtzman DM, Morris JC, Goate AM (2014) Genome-wide association study of CSF levels of 59 alzheimer’s disease candidate proteins: significant associations with proteins involved in amyloid processing and inflammation. PLoS Genet 10, e1004758.

[51] Calsolaro V, Edison P (2016) Neuroinflammation in Alzheimer’s disease: Current evidence and future directions. Alzheimers Dement 12, 719–732.

[52] Carter SF, Scholl M, Almkvist O, Wall A, Engler H, Langstrom B, Nordberg A (2012) Evidence for astrocytosis in prodromal Alzheimer disease provided by 11C-deuterium-L-deprenyl: a multitracer PET paradigm combining 11C-Pittsburgh compound B and 18F-FDG. J Nucl Med 53, 37–46.

[53] Zhang R, Miller RG, Madison C, Jin X, Honrada R, Harris W, Katz J, Forshew DA, McGrath MS (2013) Systemic immune system alterations in early stages of Alzheimer’s disease. J Neuroimmunol 256, 38–42.

[54] Jones L, Holmans PA, Hamshere ML, Harold D, Moskvina V, Ivanov D, Pocklington A, Abraham R, Hollingworth P, Sims R, Gerrish A, Pahwa JS, Jones N, Stretton A, Morgan AR, Lovestone S, Powell J, Proitsi P, Lupton MK, Brayne C, Rubinsztein DC, Gill M, Lawlor B, Lynch A, Morgan K, Brown KS, Passmore PA, Craig D, McGuinness B, Todd S, Holmes C, Mann D, Smith AD, Love S, Kehoe PG, Mead S, Fox N, Rossor M, Collinge J, Maier W, Jessen F, Schurmann B, Heun R, Kolsch H, van den Bussche H, Heuser I, Peters O, Kornhuber J, Wiltfang J, Dichgans M, Frolich L, Hampel H, Hull M, Rujescu D, Goate AM, Kauwe JS, Cruchaga C, Nowotny P, Morris JC, Mayo K, Livingston G, Bass NJ, Gurling H, McQuillin A, Gwilliam R, Deloukas P, Al-Chalabi A, Shaw CE, Singleton AB, Guerreiro R, Muhleisen TW, Nothen MM, Moebus S, Jockel KH, Klopp N, Wichmann HE, Ruther E, Carrasquillo MM, Pankratz VS, Younkin SG, Hardy J, O’Donovan MC, Owen MJ, Williams J (2010) Genetic evidence implicates the immune system and cholesterol metabolism in the aetiology of Alzheimer’s disease. PLoS One 5, e13950.

[55] McQuade A, Blurton-Jones M (2019) Microglia in Alzheimer’s Disease: Exploring How Genetics and Phenotype Influence Risk. J Mol Biol 431, 1805–1817.

[56] Villegas-Llerena C, Phillips A, Garcia-Reitboeck P, Hardy J, Pocock JM (2016) Microglial genes regulating neuroinflammation in the progression of Alzheimer’s disease. Curr Opin Neurobiol 36, 74–81.

[57] Chin-Chan M, Navarro-Yepes J, Quintanilla-Vega B (2015) Environmental pollutants as risk factors for neurodegenerative disorders: Alzheimer and Parkinson diseases. Front Cell Neurosci 9, 124.

[58] Block ML, Calderon-Garciduenas L (2009) Air pollution: mechanisms of neuroinflammation and CNS disease. Trends Neurosci 32, 506–516.

[59] Dhaini HR (2009) Neuroinflammation: Modulating Pesticide-induced Neurodegeneration In Encyclopedia of Neuroscience, Binder MD, Hirokawa N, Windhorst U, eds. Springer Berlin Heidelberg, Berlin, Heidelberg, pp. 2734-2739.

[60] Parron T, Requena M, Hernandez AF, Alarcon R (2011) Association between environmental exposure to pesticides and neurodegenerative diseases. Toxicol Appl Pharmacol 256, 379–385.

[61] Richardson JR, Roy A, Shalat SL, von Stein RT, Hossain MM, Buckley B, Gearing M, Levey AI, German DC (2014) Elevated serum pesticide levels and risk for Alzheimer disease. JAMA Neurol 71, 284–290.

[62] Fleming L, Mann JB, Bean J, Briggle T, Sanchez-Ramos JR (1994) Parkinson’s disease and brain levels of organochlorine pesticides. Ann Neurol 36, 100–103.

[63] Richardson JR, Shalat SL, Buckley B, Winnik B, O’Suilleabhain P, Diaz-Arrastia R, Reisch J, German DC (2009) Elevated serum pesticide levels and risk of Parkinson disease. Arch Neurol 66, 870–875.

[64] Singh N, Chhillar N, Banerjee B, Bala K, Basu M, Mustafa M (2013) Organochlorine pesticide levels and risk of Alzheimer’s disease in north Indian population. Hum Exp Toxicol 32, 24–30.

[65] Martin TJ, Whalen MM (2017) Exposures to the environmental toxicants pentachlorophenol (PCP) and dichlorodiphenyltrichloroethane (DDT) modify secretion of interleukin 1-beta (IL-1beta) from human immune cells. Arch Toxicol 91, 1795–1808.

[66] Martin TJ, Maise J, Gabure S, Whalen MM (2019) Exposures to the environmental contaminants pentachlorophenol and dichlorodiphenyltrichloroethane increase production of the proinflammatory cytokine, interleukin-1beta, in human immune cells. J Appl Toxicol.

[67] Massawe R, Drabo L, Whalen M (2017) Effects of pentachlorophenol and dichlorodiphenyltrichloroethane on secretion of interferon gamma (IFNgamma) and tumor necrosis factor alpha (TNFalpha) from human immune cells. Toxicol Mech Methods 27, 223–235.

[68] Cardenas-Gonzalez M, Gaspar-Ramirez O, Perez-Vazquez FJ, Alegria-Torres JA, Gonzalez-Amaro R, Perez-Maldonado IN (2013) p,p’-DDE, a DDT metabolite, induces proinflammatory molecules in human peripheral blood mononuclear cells "in vitro". Exp Toxicol Pathol 65, 661–665.

[69] Narahashi T (1992) Nerve membrane Na+ channels as targets of insecticides. Trends Pharmacol Sci 13, 236–241.

[70] Narahashi T (1996) Neuronal ion channels as the target sites of insecticides. Pharmacol Toxicol 79, 1–14.

[71] Narahashi T, Haas HG (1967) DDT: interaction with nerve membrane conductance changes. Science 157, 1438–1440.

[72] Liang CP, Sack C, McGrath S, Cao Y, Thompson CJ, Robin LP (2021) US Food and Drug Administration regulatory pesticide residue monitoring of human foods: 2009-2017. Food Addit Contam Part A Chem Anal Control Expo Risk Assess 38, 1520–1538.

[73] (CDC) CfDCaP (2019) Fourth National Report on Human Exposure to Environmental Chemicals. 1.

[74] (CDC) CfDCaP (2019) Fourth National Report on Human Exposure to Environmental Chemicals 2.

[75] Koureas M, Rousou X, Haftiki H, Mouchtouri VA, Rachiotis G, Rakitski V, Tsakalof A, Hadjichristodoulou C (2019) Spatial and temporal distribution of p,p’-DDE (1dichloro2,2bis (pchlorophenyl) ethylene) blood levels across the globe. A systematic review and meta-analysis. Sci Total Environ 686, 440-451.

[76] Silver KS, Du Y, Nomura Y, Oliveira EE, Salgado VL, Zhorov BS, Dong K (2014) Voltage-Gated Sodium Channels as Insecticide Targets. Adv In Insect Phys 46, 389–433.

[77] Roy ML, Narahashi T (1992) Differential properties of tetrodotoxin-sensitive and tetrodotoxin-resistant sodium channels in rat dorsal root ganglion neurons. J Neurosci 12, 2104–2111.

[78] Black JA, Westenbroek R, Minturn JE, Ransom BR, Catterall WA, Waxman SG (1995) Isoform-specific expression of sodium channels in astrocytes in vitro: immunocytochemical observations. Glia 14, 133–144.

[79] Reese KA, Caldwell JH (1999) Immunocytochemical localization of NaCh6 in cultured spinal cord astrocytes. Glia 26, 92–96.

[80] Janelidze S, Stomrud E, Palmqvist S, Zetterberg H, van Westen D, Jeromin A, Song L, Hanlon D, Tan Hehir CA, Baker D, Blennow K, Hansson O (2016) Plasma beta-amyloid in Alzheimer’s disease and vascular disease. Sci Rep 6, 26801.

[81] Grubman A, Choo XY, Chew G, Ouyang JF, Sun G, Croft NP, Rossello FJ, Simmons R, Buckberry S, Landin DV, Pflueger J, Vandekolk TH, Abay Z, Zhou Y, Liu X, Chen J, Larcombe M, Haynes JM, McLean C, Williams S, Chai SY, Wilson T, Lister R, Pouton CW, Purcell AW, Rackham OJL, Petretto E, Polo JM (2021) Transcriptional signature in microglia associated with Abeta plaque phagocytosis. Nat Commun 12, 3015.

[82] Kim RE, Shin CY, Han SH, Kwon KJ (2020) Astaxanthin Suppresses PM2.5-Induced Neuroinflammation by Regulating Akt Phosphorylation in BV-2 Microglial Cells. Int J Mol Sci 21.

[83] Greve HJ, Mumaw CL, Messenger EJ, Kodavanti PRS, Royland JL, Kodavanti UP, Block ML (2020) Diesel exhaust impairs TREM2 to dysregulate neuroinflammation. J Neuroinflammation 17, 351.

[84] Sayed FA, Kodama L, Fan L, Carling GK, Udeochu JC, Le D, Li Q, Zhou L, Wong MY, Horowitz R, Ye P, Mathys H, Wang M, Niu X, Mazutis L, Jiang X, Wang X, Gao F, Brendel M, Telpoukhovskaia M, Tracy TE, Frost G, Zhou Y, Li Y, Qiu Y, Cheng Z, Yu G, Hardy J, Coppola G, Wang F, DeTure MA, Zhang B, Xie L, Trajnowski JQ, Lee VMY, Gong S, Sinha SC, Dickson DW, Luo W, Gan L (2021) AD-linked R47H-TREM2 mutation induces disease-enhancing microglial states via AKT hyperactivation. Sci Transl Med 13, eabe3947.

[85] Ocanas SR, Pham KD, Cox JEJ, Keck AW, Ko S, Ampadu FA, Porter HL, Ansere VA, Kulpa A, Kellogg CM, Machalinski AH, Thomas MA, Wright Z, Chucair-Elliott AJ, Freeman WM (2023) Microglial senescence contributes to female-biased neuroinflammation in the aging mouse hippocampus: implications for Alzheimer’s disease. J Neuroinflammation 20, 188.

[86] Sala Frigerio C, Wolfs L, Fattorelli N, Thrupp N, Voytyuk I, Schmidt I, Mancuso R, Chen WT, Woodbury ME, Srivastava G, Moller T, Hudry E, Das S, Saido T, Karran E, Hyman B, Perry VH, Fiers M, De Strooper B (2019) The Major Risk Factors for Alzheimer’s Disease: Age, Sex, and Genes Modulate the Microglia Response to Abeta Plaques. Cell Rep 27, 1293–1306 e1296.

[87] Guillot-Sestier MV, Araiz AR, Mela V, Gaban AS, O’Neill E, Joshi L, Chouchani ET, Mills EL, Lynch MA (2021) Microglial metabolism is a pivotal factor in sexual dimorphism in Alzheimer’s disease. Commun Biol 4, 711.

[88] Mathys H, Davila-Velderrain J, Peng Z, Gao F, Mohammadi S, Young JZ, Menon M, He L, Abdurrob F, Jiang X, Martorell AJ, Ransohoff RM, Hafler BP, Bennett DA, Kellis M, Tsai LH (2019) Single-cell transcriptomic analysis of Alzheimer’s disease. Nature 570, 332–337.

[89] Bonham LW, Sirkis DW, Yokoyama JS (2019) The Transcriptional Landscape of Microglial Genes in Aging and Neurodegenerative Disease. Front Immunol 10, 1170.

[90] Zhao S, Umpierre AD, Wu LJ (2024) Tuning neural circuits and behaviors by microglia in the adult brain. Trends Neurosci 47, 181–194.

