## Supplementary figures and images for "Effects of Pesticide Exposure on Neuroinflammation and Microglial Gene Expression: Relevance to Mechanisms of Alzheimer’s Disease Risk"

### Figure S1

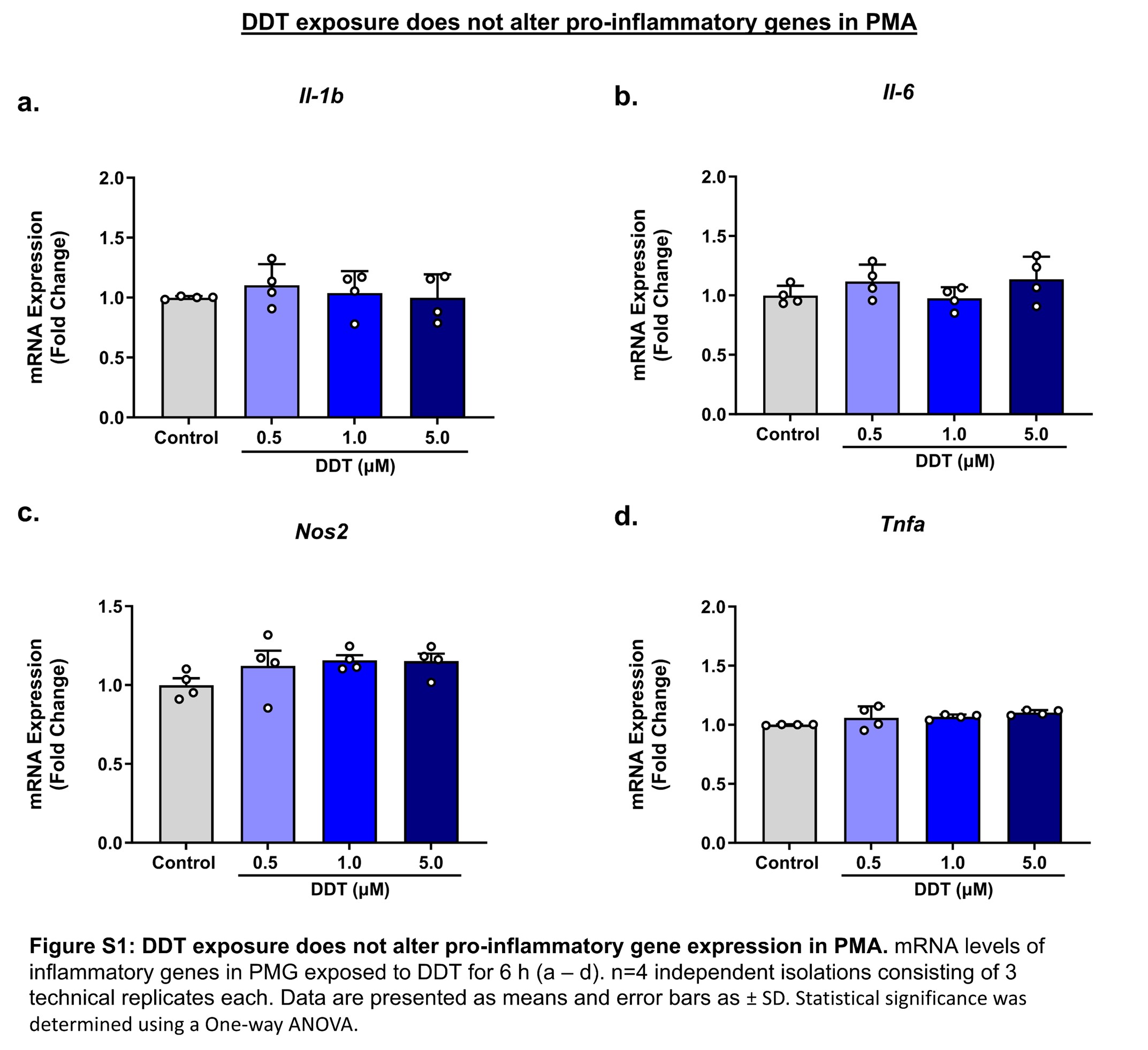
