## Supplemental Table 1 for "Effects of Pesticide Exposure on Neuroinflammation and Microglial Gene Expression: Relevance to Mechanisms of Alzheimer’s Disease Risk"

### Supplemental Table 1: Primer sequences for genes analyzed by RT-PCR

| *Il-1b* | Forward | TGGACCTTCCAGGATGAGGACA |
| --- | --- | --- |
|  | Reverse | GTTCATCTCGGAGCCTGTAGTG |
| *Il-6* | Forward | TCCATCCAGTTGCCTTCTTG |
|  | Reverse | ATTGCCATTGCACAACTCTTTT |
| *Tnfa* | Forward | GATCGGTCCCCAAAGGGATG |
|  | Reverse | CCACTTGGTGGTTTGTGAGTG |
| *Nos2* | Forward | GAGACAGGGAAGTCTGAAGCAC |
|  | Reverse | CCAGCAGTAGTTGCTCCTCTTC |
| *Cx3cr1* | Forward | GAGCATCACTGACATCTACCTCC |
|  | Reverse | AGAAGGCAGTCGTGAGCTTGCA |
| *P2ry12* | Forward | CATTGACCGCTACCTGAAGACC |
|  | Reverse | GCCTCCTGTTGGTGAGAATCATG |
| *Tmem119* | Forward | ACTACCCATCCTCGTTCCCTGA |
|  | Reverse | TAGCAGCCAGAATGTCAGCCTG |
| *Ctsb* | Forward | ACCTGTAACTGCTCACACCTC |
|  | Reverse | CTAAGAAGATGGCAGTCCGGG |
| *Tyrobp* | Forward | GAGTGACACTTTCCCAAGATGC |
|  | Reverse | CCTTGACCTCGGGAGACCA |
| *Apoe* | Forward | AGAACAACCCGCCTCGTGACA |
|  | Reverse | ATTGGCCAGTCAGCTCCTTCCG |
| *Trem2* | Forward | CTACCAGTGTCAGAGTCTCCGA |
|  | Reverse | CCTCGAAACTCGATGACTCCTC |
